# Proteomic Analysis of Urine from California Sea Lions (*Zalophus californianus*): a Resource for Urinary Biomarker Discovery

**DOI:** 10.1101/336867

**Authors:** Benjamin A. Neely, Katherine C. Prager, Alison M. Bland, Christine Fontaine, Frances M. Gulland, Michael G. Janech

**Author notes:** **Corresponding Author:** Dr. Michael G. Janech, Hollings Marine Laboratory, College of Charleston, 331 Fort Johnson Road, Charleston, SC 29412, USA.

## Abstract

Urinary markers for the assessment of kidney diseases in wild animals are limited, in part, due to the lack of urinary proteome data, especially for marine mammals. One of the most prevalent kidney diseases in marine mammals is caused by *Leptospira interrogans*, which is the second most common etiology linked to stranding of California sea lions (*Zalophus californianus*). Urine proteins from eleven sea lions with leptospirosis kidney disease and eight sea lions without leptospirosis or kidney disease were analyzed using shotgun proteomics. In total, 2694 protein groups were identified and 316 were differentially abundant between groups. Major urine proteins in sea lions were similar to major urine proteins in dogs and humans except for the preponderance of resistin, lysozyme C, and PDZ domain containing 1, which appear to be over-represented. Previously reported urine protein markers of kidney injury in humans and animals were also identified. Notably, neutrophil gelatinase-associated lipocalin, osteopontin, and epidermal fatty acid binding protein were elevated over 20-fold in the leptospirosis-infected sea lions. Consistent with leptospirosis infection in rodents, urinary proteins associated with the renin-angiotensin system were depressed, including neprilysin. This study represents a foundation from which to explore the clinical use of urinary protein markers in California sea lions.

**Abbreviations:**

RAS: renin-angiotensin system
lepto: leptospirosis
SCr: serum creatinine
BUN: blood urea nitrogen

## Introduction

Leptospirosis, the disease caused by a bacterial infection with pathogenic species within the genus *Leptospira*, is the second most common reason for California sea lions (*Zalophus californianus*) to strand sick and dying along the California coast ^1^. It is the most common cause of renal compromise in sea lions. Sea lions with leptospirosis typically present with elevated serum creatinine (SCr) and blood urea nitrogen (BUN), hypernatremia and hyperphosphatemia, and, on histological examination, have multifocal, severe lymphoplasmocytic interstitial or tubulointerstitial nephritis which may vary from acute to chronic, and tubular epithelial necrosis ^2-3^. However recent studies indicate that some animals experience mild or subclinical infections with *Leptospira*^4^and serum correlates of renal function may fail to detect mild injury. For example, in a study of sea lions with detectable anti-*Leptospira* antibodies by microscopic agglutination testing (MAT), an indication of prior exposure and possibly current infection, elevated SCr or BUN was reported for only 57.6 % and 51.5 % of seropositive animals, respectively^3^. It remains unknown whether the animals without elevated values had recovered from initial renal compromise, never experienced compromise, or were experiencing renal compromise that was not being detected by these historical correlates of renal function. In a more recent study of leptospirosis infection in sea lions, both SCr and BUN have been shown to decline to normal range even though leptospires continued to be shed in the urine ^4^, and is consistent with the idea of maintenance hosts (chronic infection without clinical indicators of disease) ^5-6^.

The utility of SCr and BUN as filtration markers is known to be imperfect and cannot differentiate pre-renal azotemia from tubular injury. Serum creatinine changes can often lag behind renal injury by up to 48 h and in the chronic setting, secretion of creatinine or reduction in creatinine production can lead to the overestimation of renal function ^7-13^. Additionally, assessment of kidney damage in surveys of sea lions with no clinical history complicates diagnosis of mild renal injury because markers such as creatinine can vary with feeding status, hydration status, muscle mass, and assay bias in the presence of confounders such as bilirubin (as reviewed in ^11^). Furthermore, reference ranges based on age class and body mass can be difficult to validate with traditional renal clearance techniques, especially when sea lions undergo peripheral vasoconstriction in response to anesthesia^14^.

Assessment of renal injury using urine protein markers has been widely described for dogs, cats, livestock and humans^15-16^, with markers including urinary total protein, albumin, immunoglobulins, as well as more recent markers of tubular injury such as neutrophil gelatinase-associated lipocalin (NGAL; approved name lipocalin 2; approved symbol LCN2). To date urine protein markers have yet to be translated to marine mammal veterinary medicine for the monitoring of renal injury. In fact, it is unclear which proteins are excreted in the urine of marine mammals and whether kidney injury markers used to monitor kidney health in domestic animals are applicable to marine mammals. The purpose of this study was to explore the urinary proteome in wild California sea lions and to compare differences in the urinary proteome between sea lions with and without leptospirosis. Search methods included the use proteome databases of closely related species. These results highlight expected and potentially novel candidate urine protein markers of tubular interstitial nephritis across a spectrum of infection and may be useful in diagnosing more subtle renal disease in sea lions, regardless of the cause.

## Experimental Section

### Sea lions

Urine samples were collected from two groups of sea lions: sea lions (Lepto, n=11) that stranded along the California coast between 2011 and 2012 and were being treated for leptospirosis and for which *Leptospira* infection status was confirmed by detection of *Leptospira* DNA in urine or kidney tissue via PCR ^17^; and sea lions (Control, n=8) which were apparently healthy, free-ranging sea lions sampled along the coasts of California (n=3) and Oregon (n=5), and which had no evidence of current (*i.e.*, negative urine *Leptopira* PCR results) or prior (*i.e.*, no detectable anti-*Leptospira* antibodies via serum MAT) ^18^ *Leptospira* infection and no evidence of renal compromise via serum chemistry analysis. Samples from Lepto animals were collected under The Marine Mammal Center (TMMC; Sausalito, CA) National Oceanic Atmospheric Administration (NOAA)/National Marine Fisheries Service (NMFS) Stranding Agreement during the course of their routine clinical care and treatment at TMMC and from Control animals during the course of a study investigating *Leptospira* exposure in free-ranging sea lions (unpublished data) under NMFS Permit No. 17115-03 and NMFS Permit No. 16087. Serum samples were collected one to five days from admission to TMMC from Lepto animals and within 48 h of urine collection. Serum and urine were collected at the same time from Control animals. Urine was collected aseptically via free catch (Lepto n=1), cystocentesis (Lepto n=1; Control n=2) or urethral catheterization (Lepto n=5; Control n=6) (Table S5) while under isoflurane anesthesia, or via needle aspiration through the wall of the bladder during necropsy (Lepto n=4). Different urine collection methods were allowed to facilitate case-control matching of animals (Table 1). Urine was stored in 2 mL cryovials at −20 to −80 °C until further processing and analysis. Serum chemistry analyses of Lepto and Control serum samples were performed on an ACE^®^ Clinical Chemistry System (Alfa Wassermann, Inc., West Caldwell, New Jersey, USA). Urine osmolality was determined using a Wescorp (Model 5100) vapor pressure osmometer. Total protein was determined using a microBCA assay. Urine creatinine was measured using a modified Jaffe method.

**Table 1.** Age, sex, and serum and urine chemistry for California sea lions in this study. Statistical probabilities were calculated using Chi-square or Mann-Whitney U test.

|  |  | Control | Lepto | p-value |
| --- | --- | --- | --- | --- |
| Sex (Male/Female) |  | 7/1 | 10/1 | *0.81* |
| Age Class |  |  |  |  |
|  | Yearling | 1 | 1 | *0.69* |
|  | Juvenile | 2 | 6 |  |
|  | Subadult | 4 | 3 |  |
|  | Adult | 1 | 1 |  |
| Serum Creatinine (SCr), mg/dL |  |  |  |  |
| | Mean ± SD | 1.1 ± 0.4 (N=8) | 5.4 ± 2.3 (N=11) | *<0.001* |
|  | Range | 0.5 - 1.6 | 2.0 - 8.9 |  |
| BUN, mg/dL |  |  |  |  |
| | Mean ± SD | 18 ± 6 (N=8) | 293 ± 181 (N=11) | *<0.001* |
|  | Range | 9 - 26 | 129 - 804 |  |
| BUN/SCr |  |  |  |  |
| | Mean ± SD | 17.5 ± 5.8 (N=8) | 60.8 ± 32.7 (N=11) | *<0.001* |
|  | Range | 12.0 - 30.2 | 21.8 - 118.2 |  |
| Plasma Na, mEq/L |  |  |  |  |
| | Mean ± SD | 149 ± 3 (N=8) | 183 ± 18 (N=10) | *<0.01* |
|  | Range | 146 - 152 | 149 - 214 |  |
| Plasma Cl, mEq/L |  |  |  |  |
| | Mean ± SD | 108 ± 3 (N=8) | 146 ± 13 (N=9) | *<0.001* |
|  | Range | 102 - 111 | 124 - 166 |  |
| Plasma Ca, mg/dL |  |  |  |  |
| | Mean ± SD | 9.6 ± 0.4 (N=8) | 8.6 ± 1.0 (N=11) | *<0.05* |
|  | Range | 9.0 - 10.1 | 7.3 - 10.2 |  |
| Plasma Phos, mg/dL |  |  |  |  |
| | Mean ± SD | 5 ± 1 (N=8) | 15 ± 9 (N=11) | *<0.001* |
|  | Range | 3 - 7 | 7 - 37 |  |
| Plasma K, mEq/L |  |  |  |  |
| | Mean ± SD | 4.4 ± 0.4 (N=8) | 4.8 ± 3.9 (N=10) | *<0.05* |
|  | Range | 3.9 - 5.1 | 2.9 - 15.8 |  |
| Urine Specific Gravity |  |  |  |  |
| | Mean ± SD | 1.034 ± 0.011 (N=8) | 1.020 ± 0.002 (N=9) | *<0.05* |
|  | Range | 1.010 - 1.045 | 1.019 - 1.021 |  |
| Urine Osmolality, mOsm/kg |  |  |  |  |
| | Mean ± SD | 1214 ± 454 (N=8) | 544 ± 83 (N=10) | *<0.01* |
|  | Range | 296 - 1577 | 420 - 661 |  |
| Urine Creatinine , mg/dL |  |  |  |  |
| | Mean ± SD | 266 ± 175 (N=8) | 57 ± 35 (N=11) | *<0.01* |
|  | Range | 72 - 513 | 20 - 136 |  |
| Urine Creatinine / Serum Creatinine |  |  |  |  |
| | Mean ± SD | 249 ± 137 (N=8) | 11 ± 5 (N=11) | *<0.001* |
|  | Range | 60 - 516 | 4 - 21 |  |
| Urine Protein, mg/dL |  |  |  |  |
| | Mean ± SD | 39.0 ± 25.4 (N=8) | 583.5 ± 1097.9 (N=11) | *0.001* |
|  | Range | 12.7 - 83.8 | 56.6 - 3503.3 |  |
| Urine Protein/Urine Creatinine |  |  |  |  |
| | Mean ± SD | 0.26 ± 0.3 (N=8) | 19.05 ± 42.4 (N=11) | *<0.001* |
|  | Range | 0.03 - 0.94 | 0.58 - 139.85 |  |

### Trypsin digestion

Urines were thawed rapidly at 37 °C in a water bath and vortexed for 5 min prior to centrifugation at 3000 x *g*_n_ for 30 s. Urine protein concentration was determined using the pyrogallol method (QuanTtest^®^ Red, Total Protein Assay System, Quantimetrix Corp.). In Lo-Bind microcentrifuge tubes (Eppendorf), urine supernatant equal to 20 μg total protein was brought to a volume of 175 μL and spiked with 400 ng HIV GP80 as an internal standard to assess digestion. Proteins were sequentially digested with proteases LysC and Trypsin (Promega) following in-solution protocols with sodium deoxycholate as a detergent previously published ^19^. Briefly, sodium deoxycholate was added to the urine protein solution to a final volume fraction of 5 %. Proteins were reduced with dithiotretitol (final concentration 10 mmol/L) for 30 min at 60 °C and subsequently alkylated with iodoacetamide (final concentration 15 mmol/L) at room temperature in the dark for 30 min. Samples were diluted eight times by volume with 50 mmol/L ammonium bicarbonate, followed by addition of LysC protease at a 1:100 mass ratio (LysC:total protein), and the samples were incubated at 37 °C for 2 h. Trypsin was then added at a 1:20 mass ratio (trypsin:total protein) and incubated overnight at 37 °C. Trifluoroacetic acid (10 % volume fraction) was added to the sample to achieve a final volume fraction of 5% trifluoroacetic acid. The tryptic peptide solution underwent liquid:liquid extraction with water-saturated ethyl acetate and was repeated four times. The peptide containing bottom layer was dried under vacuum to reduce the final volume to approximately 200 μL in a Speedvac. One mL of 0.1 % formic acid (volume fraction) was added to the peptide solution, and peptides were extracted using solid phase cartridges (Strata-x, Phenomenex), then eluted in 40 % acetonitrile (volume fraction) in 0.1 % formic acid (volume fraction). Peptides were freeze-dried under vacuum and resuspended in 0.1 % formic acid (volume fraction) prior to separation and data acquisition by nanoLC-MS/MS.

### Shotgun Proteomics (LC-MS/MS)

Peptide mixtures (approximately 1 μg) were analyzed using an UltiMate 3000 Nano LC coupled to a Fusion Lumos mass spectrometer (Thermo Fisher Scientific). Prior to analysis the sample order was randomized. Resulting peptide mixtures (10 μL) were loaded onto a PepMap 100 C18 trap column (75 μm id x 2 cm length; Thermo Fisher Scientific) at 3 μL/min for 10 min with 2 % acetonitrile (volume fraction) and 0.05% trifluoroacetic acid (volume fraction) followed by separation on an Acclaim PepMap RSLC 2 μm C18 column (75μm id x 25 cm length; Thermo Fisher Scientific) at 40°C. Peptides were separated along a 130 min gradient of 5 % to 27.5 % mobile phase B [80 % acetonitrile (volume fraction), 0.08 % formic acid (volume fraction)] over 105 min followed by a ramp to 40 % mobile phase B over 15 min and lastly to 95 % mobile phase B over 10 min at a flow rate of 300 nL/min. The Fusion Lumos was operated in positive polarity and data dependent mode (topN, 3 s cycle time) with a dynamic exclusion of 60 s (with 10 ppm error). Full scan resolution using the orbitrap was set at 120 000 and the mass range was set to 375 to 1500 *m/z*. Full scan ion target value was 4.0×10^5^ allowing a maximum injection time of 50 ms. Monoisotopic peak determination was used, specifying peptides and an intensity threshold of 1.0×10^4^ was used for precursor selection. Data-dependent fragmentation was performed using higher-energy collisional dissociation (HCD) at a normalized collision energy of 32 with quadrupole isolation at 0.7 *m/z* width. The fragment scan resolution using the orbitrap was set at 30 000, 110 *m/z* as the first mass, ion target value of 2.0×10^5^and 60 ms maximum injection time.

Raw files were converted to peak lists in the mgf format using Proteome Discoverer (v2.0.0.802). These data were searched using the Mascot algorithm (v2.6.0; Matrix Science). The following databases specified within Mascot: *Leptonychotes weddellii* (Weddell seal), NCBI RefSeq Release 100, GCF_000349705.1_LepWed1.0, 25 718 sequences; *Odobenus rosmarus divergens* (Pacific walrus), NCBI RefSeq Release 101, GCF_000321225.1_Oros_1.0, 31 370 sequences; iRT peptide, 1 sequence; HIV envelope glycoprotein, 2 sequences (UniprotKB Q72502V and P04577V); and the common Repository of Adventitious Proteins database (cRAP; 2012.01.01; the Global Proteome Machine), 107 sequences. The following search parameters were used: trypsin was specified as the enzyme allowing for two mis-cleavages; carbamidomethyl (C) was fixed and deamidated (NQ) pyro-Glu (n-term Q), and oxidation (M) were variable modifications; 10 ppm precursor mass tolerance and 0.02 Da fragment ion tolerance; instrument type was specified as ESI-FTICR; the decoy setting was used within Mascot to provide local FDR. Mascot results were confirmed using Scaffold (v. 4.7.2; Proteome Software, Portland, OR, USA), with a minimum protein threshold of 1 % false discovery rate (FDR), and 1 % FDR at the peptide level. Experiment wide grouping was used which resulted in 2694 protein families identified at 0.9 % FDR (as determined by Mascot’s local FDR), after removal of cRAP and decoy hits. Weighted spectrum counts were exported from Scaffold for downstream analysis. In order to evaluate the differentially abundant proteins and proteins relevant to AKI and RAS pathways, these protein families were converted to a human equivalent protein with a HUGO Gene Nomenclature Committee (HGNC) gene symbol. This was based either on the NCBI RefSeq protein description exactly matching an HGNC description or on human sequence similarity search results via NCBI’s BLAST (Basic Local Alignment Search Tool). Although these names and symbols often align, there were four notable exceptions where the HGNC approved name and symbol have diverged from the common clinical name: KIM1 (kidney injury molecule 1; HGNC approved symbol HAVCR1), NGAL (neutrophil gelatinase-associated lipocalin; HGNC approved symbol LCN2), neprilysin (HGNC approved symbol MME), and osteopontin (HGNC approved symbol SPP1). Raw data, search results and databases have been deposited to the ProteomeXchange Consortium via the PRIDE^20^ partner repository with the dataset identifier PXD009019 and 10.6019/PXD009019.

### Data analysis

For all analysis unless otherwise stated, Matlab (R2015a) was used. For differential analysis of proteomic data, a two-sided Wilcoxon rank sum test was utilized. False discovery rate for multiple hypothesis testing was estimated using the method of Benjamini and Hochberg (BH; ^21^). Differentially abundant proteins were defined as a BH corrected p-value < 0.05. Log2 fold-change values were calculated as the difference in log-average counts between groups (if average counts were 0, then half of the smallest group average was used, which was 0.0427). For display purposes only, a custom implementation of a membership function was used to normalize the weighted spectral counts to a value between 0 and 1 before clustering. These normalized values were used for hierarchical clustering with a Euclidean distance metric. Differences between clinical and demographic data were determined using a Mann-Whitney U test or Chi-Square. Groups were deemed different if *p*-value < 0.05. Gene ontology and pathway inference was generated by the Reactome Pathway Database ^22^. Protein accession numbers were converted to homologous human accession numbers for ontology and pathways analysis.

## Results

### Sea lion blood and urine chemistry

Sea lions with leptospirosis kidney disease, *i.e.*, Lepto animals, displayed elevations in serum creatinine (SCr), blood urea nitrogen (BUN), plasma sodium (Na), plasma chloride (Cl), plasma potassium (K), and plasma phosphorus (P) (Table 1) compared to age and sex matched animals, characteristic of a renal disease phenotype. Measures of urine concentration (urine creatinine, urine osmolality, urine specific gravity, and urine creatinine/serum creatinine) were depressed in sea lions with leptospirosis consistent with tubular injury and a reduction in urine concentrating ability (Table 1). Renal injury in the Lepto group was also markedly depicted by the 15-fold elevation in mean urine protein concentration and 73-fold elevation in urine protein/urine creatinine ratio (Table 1). Twenty-four hour renal protein excretion was not measured.

### Sea lion urine proteins

There were 2694 protein groups identified across the analysis, excluding contaminant proteins (Table S1). Proteins identified from sea lions in the Control group represented 1636 in total with a mean and standard deviation of 794 ± 162 proteins per individual. Proteins identified in the Lepto group represented 2235 in total, with a mean and standard deviation of 831 ± 408 proteins per individual. There was no difference between the number of proteins identified between groups (*p* > 0.8; Mann-Whitney U test).

For Control sea lions the most abundant protein in the urine as determined by spectral counting was albumin, followed by uromodulin and protein AMBP. Of the 34 most abundant proteins in sea lion urine (Figure 1), several ranked proteins were unexpected compared to a rank classification of relative abundance reported for the human urine proteome ^23^. Resistin was the 11^th^ top ranked protein in sea lion urine; whereas, in humans resistin is ranked 414 ^23^. Similarly, lysozyme C and PDZ domain containing 1 were found at a much lower rank (*i.e.*, higher abundance) compared to normal human urine (human rank: 525 and 1002, respectively) and PDZ domain containing 1 was not reported in the supernatant of the dog urine proteome ^24^. Both resistin and lysozyme C belong to pathways involved in neutrophil degranulation as classified by the Reactome Knowledge Database ^22^.

**Figure 1.**
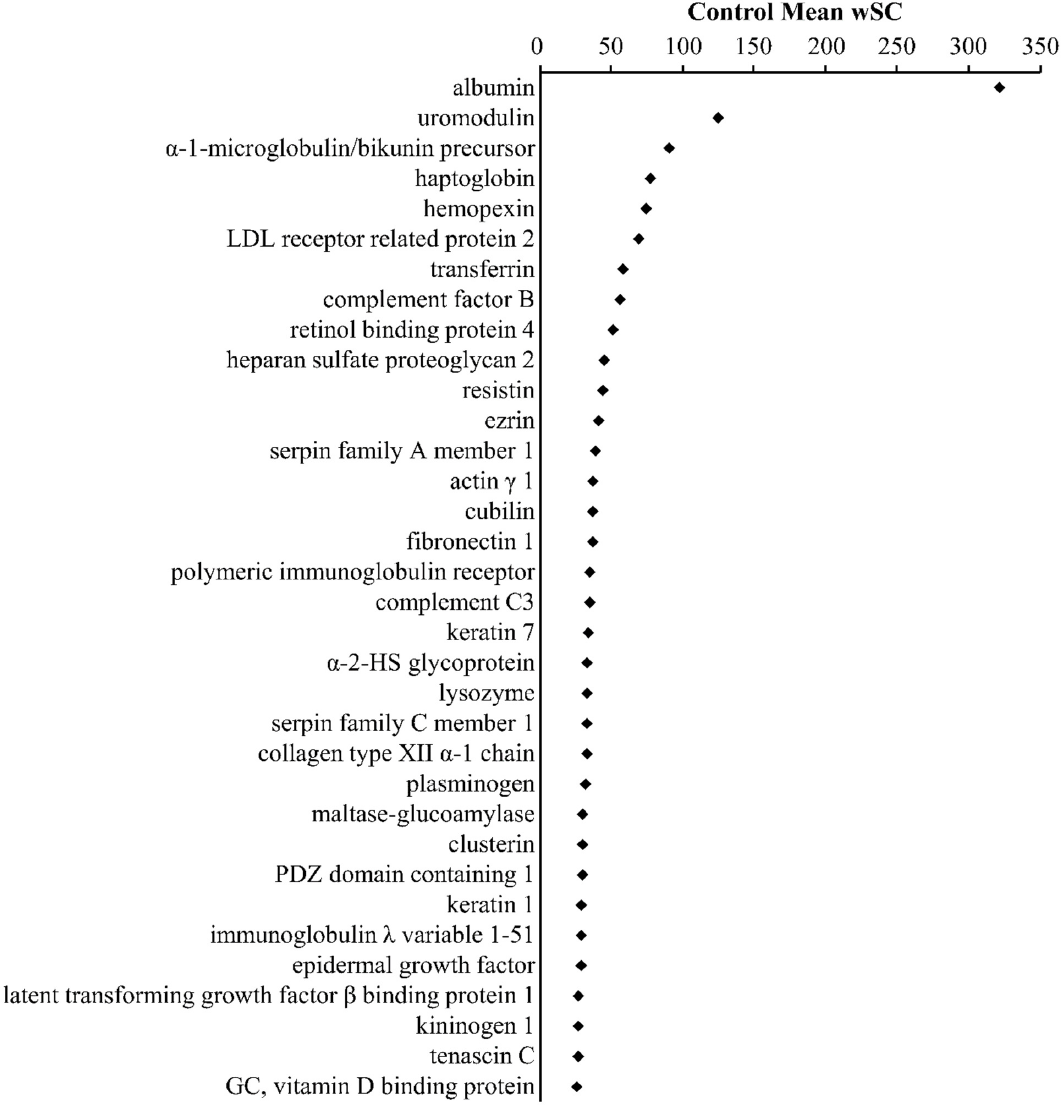
Most abundant proteins in urine from the Control group. Based on mean weighted spectral counts (wSC) in urine from the control group, there were 34 proteins that comprised 1/3 of weighted spectral counts from the urine. The top two proteins, albumin and uromodulin comprise 8.5 % of the weighted spectral counts in urine. Protein names were humanized to HGNC approved names.

### Protein differences: Control versus Lepto

There were 316 proteins differentially abundant proteins between the control and leptospirosis groups (Benjamini-Hochberg corrected *p* < 0.05; Wilcoxon rank sum test). Of these, 98 proteins were elevated and 218 proteins reduced in the Lepto group compared to the Control group. (Figure 2, Table S1). Of these differential proteins, osteopontin (HGNC name: secreted phosphoprotein 1; HGNC gene symbol, SPP1) displayed the greatest positive log_2_-fold change (log_2_ FC) relative to the Control group (log_2_ FC, 8.28), and AXL receptor tyrosine kinase (HGNC gene symbol, AXL) displayed the greatest negative log_2_-fold change (log_2_FC, −7.46). Reactome overrepresentation analysis (Table S1) of positively changing proteins (Lepto versus Control) resulted in significant grouping within pathways: 1) Platelet degranulation, 2) Complement cascade, 3) Regulation of Insulin-like Growth Factor transport and uptake by Insulin-like Growth Factor Binding Proteins. Overrepresentation analysis of negative changing proteins (Lepto versus Control) resulted in significant grouping within pathways: 1) Post-translational protein phosphorylation, 2) Regulation of Insulin-like Growth Factor transport and uptake by Insulin-like Growth Factor Binding Proteins, 3) Molecules associated with elastic fibers, 4) Neutrophil degranulation.

**Figure 2.**
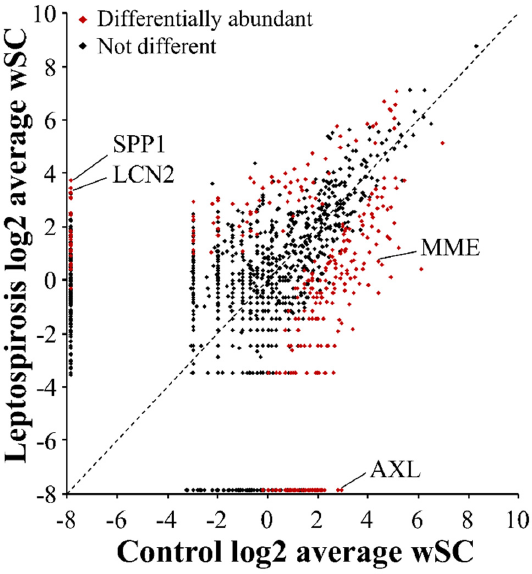
Comparison of mean weighted spectral counts between groups. There were 2694 proteins identified and the log_2_ mean weighted spectral counts (wSC) of each protein in the Control and Lepto groups were plotted. In the case the group average was zero, 0.0427 was used (half the minimum observed group average wSC). Points along the dotted line are those equal in both groups, while farther from the line indicates greater differences between groups. Of these proteins, 316 were differentially abundant (Wilcoxon rank sum, Benjamini-Hochberg adjusted *p*-value < 0.05); 98 were increased in the Lepto group and 218 were increased in the control group. Four proteins of interest are indicated: SPP1 (secreted phosphoprotein 1; alternative name osteopontin); LCN2 (lipocalin 2; alternative symbol, NGAL); AXL (AXL receptor tyrosine kinase); MME (membrane metalloendopeptidase; alternative name, neprilysin).

### Urinary protein markers of renal injury in sea lions

Select urine protein marker protein classes related to kidney injury in humans and domestic veterinary animals (primarily extracted from references within ^15-16, 25^) were clustered across samples to display the relationship of these markers with Lepto and Control sea lion groups (Figure 3 and Table S2). Notably, neutrophil gelatinase-associated lipocalin (NGAL; HGNC name, lipocalin 2; HGNC gene symbol LCN2), osteopontin (HGNC gene symbol SPP1), and epidermal-type fatty-acid binding protein (HGNC gene symbol FABP5) were elevated more than 20-fold (log_2_ FC, >4.3) over the Control group. Other previously reported markers of kidney injury that were elevated in the Lepto group include: clusterin (HGNC gene symbol CLU) and tissue inhibitor of matrix metalloproteinase 1 (HGNC symbol TIMP1).

**Figure 3.**
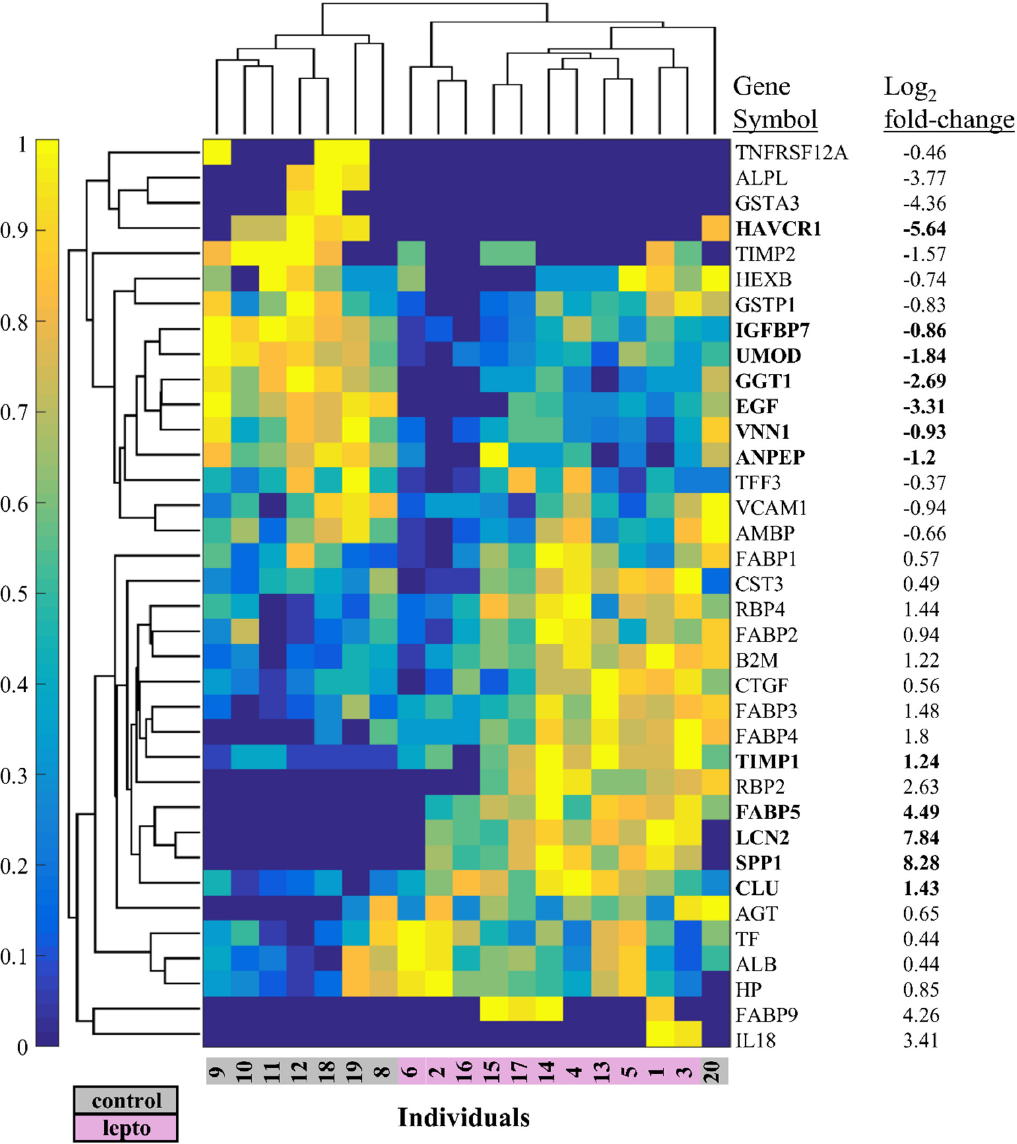
Clustering of protein markers of kidney injury reported from other mammals. Hierarchical clustering was used on 38 urine proteins detected in sea lion urine and considered as markers of kidney injury from human or domestic veterinary studies. Log_2_fold-change values are Lepto/Control. Weighted spectral counts were normalized using the membership function (normalized values are 0 to 1). Bold indicates differentially abundant proteins (Wilcoxon rank sum using weighted spectral counts, Benjamini-Hochberg adjusted *p*-value < 0.05). AGT (angiotensinogen), ALB (albumin), ALPL (alkaline phosphatase, liver/bone/kidney), AMBP (alpha-1-microglobulin/bikunin precursor), ANPEP (alanyl aminopeptidase, membrane), B2M (beta-2-microglobulin), CLU (clusterin), CST3 (cystatin C), CTGF (connective tissue growth factor), EGF (epidermal growth factor), FABP1 (fatty acid binding protein 1), FABP2 (fatty acid binding protein 2), FABP3 (fatty acid binding protein 3), FABP4 (fatty acid binding protein 4), FABP5 (fatty acid binding protein 5), FABP9 (fatty acid binding protein 9), GGT1 (gamma-glutamyltransferase 1), GSTA3 (glutathione S-transferase alpha 3), GSTP1 (glutathione S-transferase pi 1), HAVCR1 (hepatitis A virus cellular receptor 1; alternative symbol, KIM1), HEXB (hexosaminidase subunit beta), HP (haptoglobin), IGFBP7 (insulin like growth factor binding protein 7), IL18 (interleukin 18), LCN2 (lipocalin 2; alternative symbol, NGAL), RBP2 (retinol binding protein 2), RBP4 (retinol binding protein 4), SPP1 (secreted phosphoprotein 1), TF (transferrin), TFF3 (trefoil factor 3), TIMP1 (TIMP metallopeptidase inhibitor 1), TIMP2 (TIMP metallopeptidase inhibitor 2), TNFRSF12A (TNF receptor superfamily member 12A), UMOD (uromodulin), VCAM1 (vascular cell adhesion molecule 1), VNN1 (vanin 1). Values based on the walrus and Wedell sea lion database are displayed.

The following urinary markers of kidney injury were reduced in the Lepto group compared to Control: uromodulin (HGNC gene symbol UMOD), gamma-glutamyltranspeptidase 1 (HGNC gene symbol GGT1), vanin 1 (HGNC gene symbol VNN1), insulin like growth factor binding protein 7 (HGNC gene symbol IGFBP7), and alanyl aminopeptidase (HGNC gene symbol ANPEP), and kidney injury molecule 1 (KIM1; HGNC name, hepatitis A virus cellular receptor 1; HGNC gene symbol HAVCR1).

The following urinary markers of renal injury were not differentially abundant between Control and Lepto groups: interleukin 18 (HGNC gene symbol IL18), beta-2-microglobulin (HGNC gene symbol B2M), liver type fatty acid binding protein (HGNC gene symbol FABP1), transferrin (HGNC gene symbol TF), albumin (HGNC gene symbol ALB), retinal binding protein 4 (HGNC gene symbol RBP4), cystatin C (HGNC gene symbol CST3), alkaline phosphatase (HGNC gene symbol ALPL), alpha-1-microglobulin (HGNC gene symbol AMBP), haptoglobin (HGNC gene symbol HP), tissue inhibitor of matrix metalloproteinase 2 (HGNC gene symbol, TIMP2), angiotensinogen (HGNC gene symbol AGT), N-acetyl-β-D-glucosaminidase (HGNC gene symbol HEXB), alpha glutathione-S-transferase (HGNC gene symbol GSTA3), glutathione S-transferase P-like (HGNC gene symbol, GSTP1), and epidermal growth factor (HGNC gene symbol EGF).

Putative renal injury markers not found in the results include: YKL-40 (HGNC gene symbol CHI3L1), C-reactive protein (HGNC gene symbol CRP), cysteine rich protein 61 (HGNC gene symbol CYR61), sodium/hydrogen exchanger 3 (HGNC gene symbol SLC9A3).

### Renin-Angiotensin System

Based on a previous report ^26^, a reduction in polypeptide components of the renin-angiotensin system (RAS), *e.g.*, neprilysin and angiotensin I, in rodents infected with *Leptospira* ^26^appear to be characteristic of the response to infection. For this reason, data associated with these RAS proteins were specifically extracted to determine whether a similar response was characteristic of large mammals presenting with a natural course of disease (Figure 4). The following RAS-related proteins were reduced in the Lepto group: neprilysin (HGNC gene symbol MME, −3.69 log_2_ FC), angiotensin converting enzyme 1 (HGNC gene symbol ACE, −3.46 log_2_ FC), angiotensin converting enzyme 2 (HGNC gene symbol ACE2, −2.50 log_2_ FC), glutamyl aminopeptidase (alternative name, aminopeptidase A; HGNC gene symbol ENPEP, −3.36 log_2_FC), alanyl aminopeptidase (alternative name, aminopeptidase N; HGNC gene symbol ANPEP, - 1.2 log_2_ FC), carboxypeptidase A1 (HGNC gene symbol CPA1, −2.62 log_2_ FC), cathepsin D (HGNC gene symbol CTSD, −0.8 log_2_ FC). The following RAS-related proteins were not differentially abundant between groups: angiotensinogen (HGNC gene symbol AGT), carboxypeptidase A2 (HGNC gene symbol CPA2), cathepsin B (HGNC gene symbol CTSB), aspartyl aminopeptidase (HGNC gene symbol DNPEP), prolyl endopeptidase (HGNC gene symbol PREP), dipeptidyl peptidase 3 (HGNC gene symbol DPP3), cathepsin G (HGNC gene symbol CTSG), renin receptor (HGNC gene symbol ATP6AP2). The following RAS-related proteins were not identified: renin (HGNC gene symbol REN), chymase (HGNC gene symbol CMA1), thimet oligopeptidase (HGNC gene symbol THOP1), and prolyl carboxypeptidase (HGNC gene symbol PRCP).

**Figure 4.**
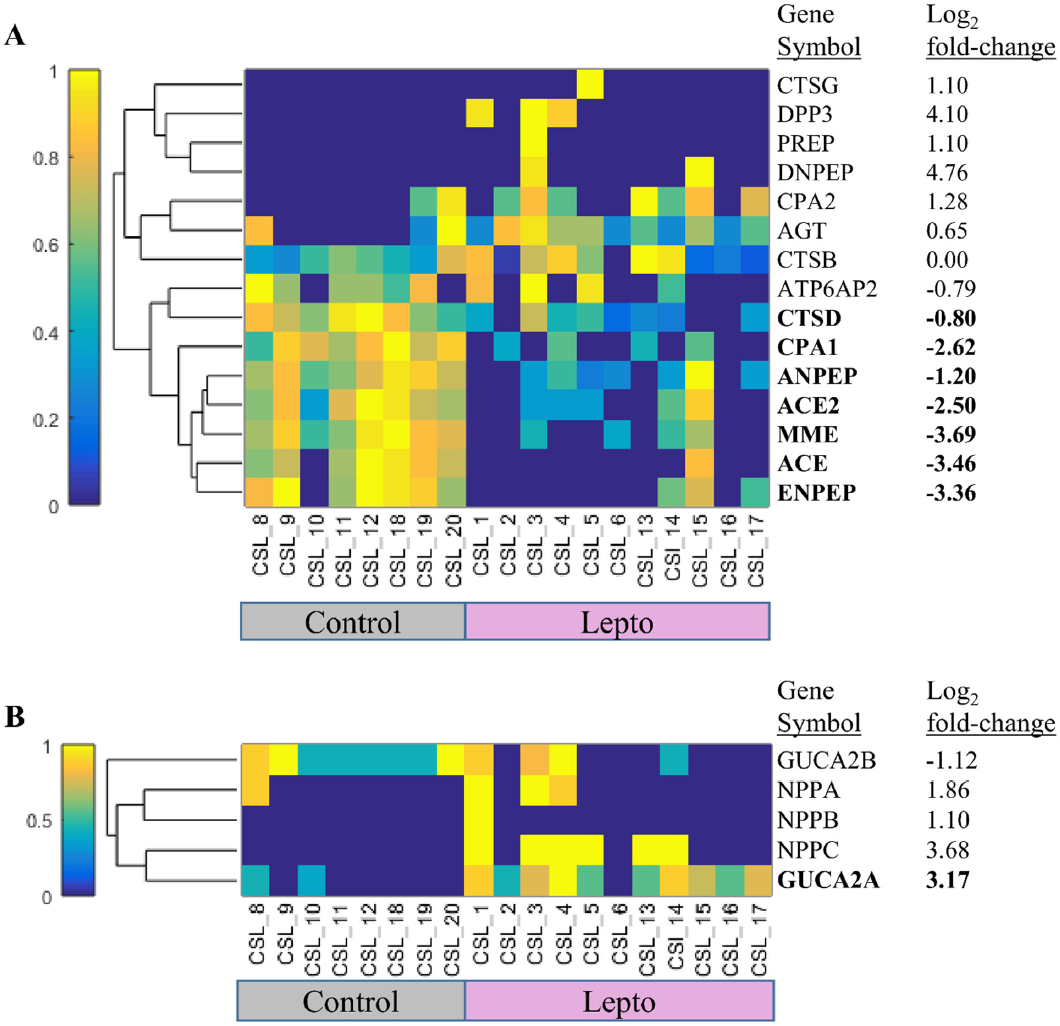
Clustering of proteins in the Renin-Angiotensin System and major Natriuretic proteins. **A.** Hierarchical clustering was used on 15 proteins in the Renin-Angiotensin System. Log2 fold-change is Lepto/Control. Weighted spectral counts were normalized using the membership function (normalized values are 0 to 1). Bold indicates differentially abundant proteins (Wilcoxon rank sum using weighted spectral counts, Benjamini-Hochberg adjusted *p*-value < 0.05). CTSG (cathepsin G), DPP3 (dipeptidyl peptidase 3), PREP (prolyl endopeptidase), DNPEP (aspartyl aminopeptidase), CPA2 (carboxypeptidase A2), AGT (angiotensinogen), CTSB (cathepsin B), ATP6AP2 (ATPase H+ transporting accessory protein 2), CTSD (cathepsin D), CPA1 (carboxypeptidase A1), ANPEP (alanyl aminopeptidase, membrane; alternative name, aminopeptidase N), ACE2 (angiotensin I converting enzyme 2), MME (membrane metalloendopeptidase; alternative name, neprilysin), ACE (angiotensin I converting enzyme), ENPEP (glutamyl aminopeptidase; alternative name, aminopeptidase A). **B.** Hierarchical clustering was used on five natriuretic peptide proteins. Log_2_fold-change is Lepto/Control. Weighted spectral counts were normalized using the membership function (normalized values are 0 to 1). Bold indicates differentially abundant proteins (Wilcoxon rank sum using weighted spectral counts, Benjamini-Hochberg adjusted *p*-value < 0.05). GUCA2B (Guanylate cyclase activator 2B), NPPA (Natriuretic peptide A), NPPB (Natriuretic peptide B), NPPC (Natriuretic peptide C), GUCA2A (Guanylate cyclase activator 2A).

The reduction in the RAS enzyme, neprilysin (HGNC gene symbol MME), was validated using Western blotting (Figure 5). A single immunoreactive protein band was detected between 75 kDa and 100 kDa (predicted molecular weight 86 kDa). Densitometry of the band across individual sea lions indicated a significant reduction (*p* < 0.01, Mann-Whitney U test) in urinary neprilysin protein normalized either by total protein loaded on the gel (Figure 5B) or normalized per volume of urine (Figure 5C).

**Figure 5.**
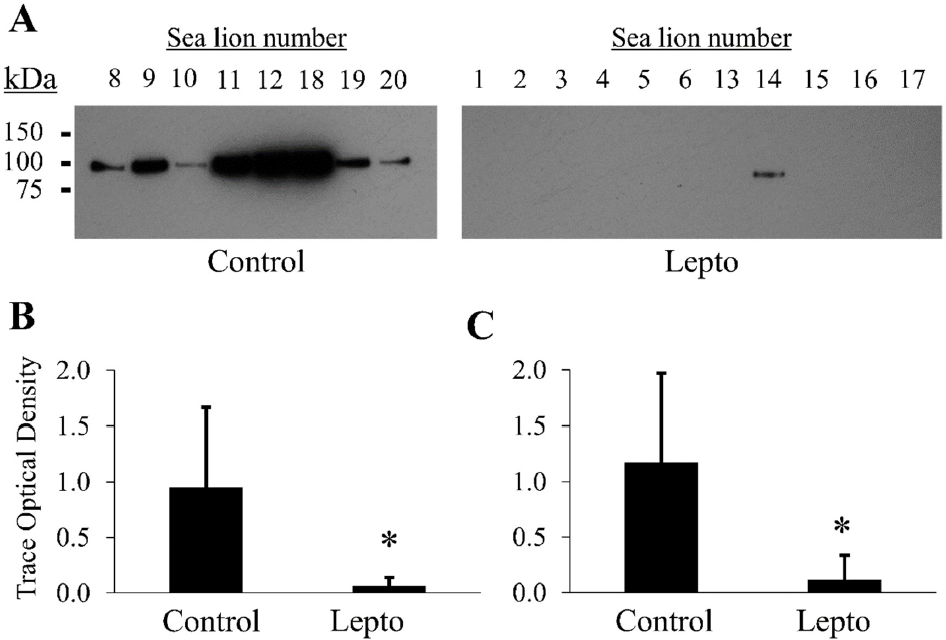
Western blot for urinary Neprilysin (HGNC gene symbol MME). Equal protein (3 μg) was loaded per lane **(A)**. Immunoreactivity was detected as a single band between 75 kDa and 100 kDa. Secondary antibody alone control blots were negative. Images were cropped and contrast modified for display purposes. Semi-quantitative comparison of urinary neprilysin immunoreactivity by densitometry normalized to total urine protein **(B)** or total volume (5 μl, **C**) loaded per lane. Asterisks indicate *p* < 0.01, Mann-Whitney U test. Error bars represent ± standard deviation.

### Natriuretic proteins

Leptospirosis associated with renal failure leads to a hypernatremic state in sea lions, therefore we extracted information regarding natriuretic proteins from the urine data for description. The following natriuretic proteins were identified: guanylin (HGNC gene symbol GUCA2A), uroguanylin (alternative name, guanylate cyclase activator 2B; HGNC gene symbol GUCA2B), C-type natriuretic peptide (HGNC gene symbol NPPC), natriuretic peptides A (HGNC gene symbol NPPA), and natriuretic peptides B (HGNC gene symbol NPPB). Only guanylin was significantly elevated in sea lions from the Lepto group.

## Discussion

California sea lions excrete a urine similar in protein composition to other mammals ^23-24,27^, where proteins such as albumin, uromodulin, and protein AMBP dominate abundance (Figure 1, TableS1). Unlike rodents, where the urine is dominated by major urinary proteins ^28^, or cats, in which the urinary protein cauxin dominates the proteome ^27^; sea lions excrete a repertoire of proteins that closely parallels that of dogs. ^24^ and humans ^23^. Some features of the sea lion urinary proteome were unexpected such as the predominance of two low molecular weight proteins, resistin (12 kDa) and lysozyme C (16 kDa), as estimated by ranking proteins by spectral counts in Control group animals. Both proteins are elevated in immune cells either in response to lipopolysaccharide to promote inflammation ^29^ or in defense of elevated lipopolysaccharide to promote bacterial lysis ^30^, respectively. Over-abundance of these two innate immune system proteins may indicate elevated innate protection against pathogenic, considering likely constant environmental pathogen exposure in the wild.

### Leptospirosis

In an effort to extend the utility of this study into veterinary diagnostics, *e.g.*, discovery of candidate urine markers of interstitial nephritis, urine proteins from sea lions with leptospirosis kidney disease were included as a separate group. Sea lions in the leptospirosis (Lepto) group included in this study displayed a range of urine protein to creatinine ratios and SCr levels that may reflect a range of renal injury severity and chronicity. Sea lions accepted for rehabilitation are discovered during strand surveys or citizen reports, therefore the clinical history, and hence the duration of infection, is unknown. Inclusion of data from the Lepto group resulted in the identification of an additional 1058 proteins, providing a more comprehensive urine proteome picture which was derived from urine from sea lions experiencing severe renal compromise as well as from those with apparently healthy kidneys. Similar to urinary proteomic results from rats with chronic *Leptospira* infection ^26^, sea lions with leptospirosis had a larger proportion of urinary proteins that were reduced in abundance compared to uninfected sea lions. Despite significant proteinuria in the Lepto group, urinary albumin concentration was not different between groups based on weighted spectral counts. This is consistent with changes in proteinuria from tubular, not glomerular, origin as has been reported from dogs with leptospirosis ^31^.

Urinary proteins as markers of renal injury and repair continue to be extensively studied and evaluated for use as human clinical diagnostics and domestic veterinary diagnostics ^15-16^. Not surprising, marine mammal veterinary diagnostics have lagged behind these other fields. To test whether previously described renal injury markers for humans and domestic veterinary species change with renal injury status, a subset of proteins were extracted from the data based on their precedent as biomarkers of renal tubular injury (Figure 3). Neutrophil gelatinase-associated lipocalin (LCN2) and osteopontin (SPP1) were notably abundant in urine from the Lepto group. Both are considered tubular injury markers of acute kidney injury ^25^ and drug-mediated nephrotoxicosis ^32^. Whereas, neutrophil gelatinase-associated lipocalin seems to promote recovery, osteopontin appears to promote monocyte infiltration and fibrosis ^33^. Regardless of mechanism, these proteins clearly have a mechanistic role in the pathogenesis of *Leptospira* infection and potential value as urinary biomarkers similar to other species.

Fatty acid binding proteins (FABP) are abundant, low molecular weight proteins that act as lipid chaperones and play diverse roles including the formation of eicosanoids in macrophages ^34^. The most widely studied FABP relevant to kidney injury markers is the liver-type FABP ^35^,which is elevated in response to acute kidney injury of multiple etiologies. In the current study, the acute kidney injury marker liver-type FABP was not different between both groups, but interestingly, epidermal-type FABP was considerably elevated in the Lepto group relative to the Control group. Epidermal FABP is not typically associated with expression in the kidney ^36^, but has been detected in human kidney and reported in the Human Protein Atlas ^37^. Of the FABPs identified, the epidermal type may be of greater interest to prospective studies of tubular injury in sea lions, or at least should be considered alongside biomarker studies of liver-type FABPs.

Other notable differentially abundant proteins with biomarker precedence include clusterin and uromodulin. In dogs with leishmaniasis, characterized by glomerular mesangial proliferation and interstitial nephritis, urinary clusterin was elevated in more advanced renal cases ^38^. Gentamicin-induced tubular necrosis also resulted in an increase in urinary clusterin in beagles, suggesting that tubular dysfunction alone can cause increased urinary levels of this protein ^39^. In the current study, the elevation in urinary clusterin in sea lions suggests a similar urinary clusterin response is conserved in sea lions experiencing renal compromise. Uromodulin is a tubular protein primarily secreted at the thick ascending limb of the kidney and has been implicated in a myriad of roles including innate immunity and transporter regulation ^40^. Reduced uromodulin excretion is consistent with tubular injury, has been implicated as a risk factor for increased urinary tract infections in humans ^41^and has been associated with severe azotemia in dogs ^42^. Lepto sea lions also displayed a lower level of uromodulin in the urine and we hypothesize that injury response mechanisms, similar to those described for humans ^40^, are conserved in sea lions.

Tyrosine kinase receptor AXL was the protein most reduced in abundance in Lepto sea lions. Its ligand, growth arrest specific protein (HGNC gene symbol GAS6) was also reduced. GAS6/AXL is known to reduce tubulo-interstitial apoptosis in chronic kidney injury models ^43^ and upregulation of this axis promotes glomerular hypertrophy in diabetic mice ^44^. AXL expression is primarily located in vascular endothelia and the proximal tubule ^45^. A survey of human kidneys with inflammatory disease provides evidence of gross upregulation in the tissue which was shown to be elevated, in part, through Angiotensin II in cell culture models ^46^. If AXL/GAS6 expression is driven by inflammatory mechanisms, then this axis should be elevated in Lepto sea lions, but in fact, both urinary AXL and GAS6 were reduced. It remains possible that the large reduction in AXL/GAS6 could simply be due to substantial tubular damage. From a different perspective, the GAS6/AXL partners are typically reduced in the normal human kidney to the point where they are barely detectable ^46^. However, in Control sea lions, urinary GAS6/AXL appears to be abundant. Given that diving results in peripheral vasoconstriction ^47^, and that the GAS6/AXL axis has been reported to be protective of ischemia reperfusion injury ^48^, we hypothesize that GAS6/AXL may play a protective role in Control sea lions and that the loss of these proteins seen in the Lepto group may reflect a loss of renal cell survival. Despite a large number of studies of the GAS6/AXL axis in humans and laboratory rodent models ^49^, studies regarding the GAS6/AXL axis in normal marine mammal physiology are non-existent and the detection of AXL/GAS6 in Control sea lions may indicate the presence of an anti-apoptotic, pro-survival protective mechanism that is affected during *Leptospira* infection in sea lions.

Although many established tubular injury markers were not significantly altered, or in some cases levels changed in the opposing direction than expected, we cannot discount these proteins as potential urinary biomarkers for sea lions based on this study. In addition to the small numbers of individuals included in this study and lack of a species-specific proteome for the identification of proteins, low abundance peptides and proteins can lead to highly variable counts. Targeted absolute measurements with an appropriate standardized assay should always be considered in future prospective biomarker studies.

### Renin-angiotensin system (RAS) and natriuretic proteins

Curiously, rats chronically infected with *Leptospira* display a less diverse urinary pattern of polypeptides including a reduction in RAS components; neprilysin (MME; alternative name, neutral endopeptidase) and angiotensin I ^26^. The reduction in urinary neprilysin protein was again reported by 2D-DIGE analysis in rats ^50^ as well as aminopeptidase N (alternative name, aminopeptidase M; HGNC approved name alanyl aminopeptidase, membrane), both of which play a role in the metabolism of angiotensin peptides in the intrarenal RAS ^51^. Commensurate to the rat studies, sea lions with leptospirosis displayed a reduction in many of the same components of the urinary RAS including neprilysin, aminopeptidase N, aminopeptidase A (HGNC approved name glutamyl aminopeptidase), angiotensin converting enzyme, angiotensin converting enzyme 2, carboxypeptidase A, and cathepsin D. The fact that renin was not discovered in the proteomic data for any sea lion, suggests that cathepsin D may be a major player in the sea lion intrarenal RAS as cathepsin D has been shown to be able to generate angiotensin I peptide from angiotensinogen ^52^.

The reduction in neprilysin was validated by immunoblot of sea lion urine proteins normalized to total protein or urine volume. Urinary neprilysin derived from proximal tubule has been recently reported to increase in humans with acute kidney injury patients due to sepsis ^53^, which is opposite of the urinary neprilysin phenotype of rats and sea lions with leptospirosis. A decrease in urinary neprilysin has been reported in *db*/*db* diabetic mice with hyperglycemia and microalbuminuria ^54^, which may be a response common to chronic renal inflammation ^55^. The consistent finding of urinary neprilysin reduction in two very different mammalian species suggests a possible targeted role for this enzyme in the response to leptospirosis.

Two notable clinic features of sea lions with leptospirosis are hypernatremia concomitant with a low urine osmolality. Hypernatremia may present in animals with reduced access to free water, diuresis due to azotemia, and renal impairment that results in a reduced ability to concentrate urine. More specifically, nephrogenic diabetes insipidus and renal vasopressin resistance appears to contribute to the urinary concentrating defect during leptospirosis infection, which might explain the hypernatremia concomitant with low urine osmolality^56-58^. In defense against hypernatremia, humans respond by inducing a natriuresis independent of blood pressure, atrial natriuretic peptide, and the RAS ^59^. To assess whether other natriuretic protein hormones may play a role in defending against hypernatremia, a survey of urinary natriuretic proteins in the two sea lion groups implicated the natriuretic pro-hormone guanylin as a potential regulator of sodium excretion. Pro-guanylin is primarily expressed in the gasterointestinal tract, but can be freely filtered by the kidney. The finding of elevated pro-guanylin abundance was unexpected because uroguanylin is anticipated to be more relevant in terms of expression in the kidney and detectability in the urine ^60^; however, pro-uroguanylin protein did not change suggesting that intestinal or renal guanylin may be upregulated in defense of normonatremia in sea lions. The link between sodium balance and natriuresis via an intestinal-renal natriuretic axis in healthy humans has been disputed ^61^making this observation in sea lions a potentially fascinating area of discovery regarding the regulation of sodium balance in leptospirosis kidney disease and in marine mammals in general.

## Conclusions

Urine proteomes have become repositories for biomarker discovery and highlight mechanistic descriptions of normal and pathophysiology. To date, information regarding the urine proteome of marine mammals, such as the California sea lion, has been limited by a lack of proteomic efforts. There is a need for renal injury markers for marine mammals, in addition to SCr and BUN, in order to assess renal insufficiency beyond imperfect markers of filtration. Based on this study, there are many parallels between urine proteins from sea lions and published domestic veterinary and human clinical protein markers that can be used to accelerate biomarker assessment to classify mild and moderate renal injury.

## Acknowledgements

We would like to thank the veterinarians, biologists, and volunteers of The Marine Mammal Center (Sausalito, CA) and the Marine Mammal Laboratory (NOAA/NMFS) for their assistance in collecting samples. This work was supported by the National Science Foundation (OCE-1335657), the US Department of Defense Strategic Environmental Research and Development Program (RC01-020/RC-2635), the John H. Prescott Marine Mammal Rescue Assistance Grant Program, and the Hellman Family Foundation. Identification of certain commercial equipment, instruments, software or materials does not imply recommendation or endorsement by the National Institute of Standards and Technology, nor does it imply that the products identified are necessarily the best available for the purpose.

The authors declare no conflicts of interest.

## Supporting Information

Supplemental Table S1. Protein identifications and differential analysis between groups. Supplemental Table S2. Identification and weighted spectral counts of 36 reported markers of kidney injury.

Supplemental Table S3. Identification and weighted spectral counts of proteins involved in the RAS pathway.

Supplemental Table S4. Identification and weighted spectral counts of proteins involved in the natriuretic pathway.

Supplemental Table 5. Serum and urine chemistry values for individual sea lions utilized to construct Table 1.

## References

1. Greig, D. J.; Gulland, F. M. D.; Kreuder, C., A Decade of Live California Sea Lion (Zalophus californianus) Strandings Along the Central California Coast: Causes and Trends, 1991-2000. Aquatic Mammals 2005, 31 (1), 11–22.

2. Gulland, F. M.; Koski, M.; Lowenstine, L. J.; Colagross, A.; Morgan, L.; Spraker, T., Leptospirosis in California sea lions (Zalophus californianus) stranded along the central California coast, 1981-1994. J Wildl Dis 1996, 32 (4), 572–80.

3. Colagross-Schouten, A. M.; Mazet, J. A.; Gulland, F. M.; Miller, M. A.; Hietala, S., Diagnosis and seroprevalence of leptospirosis in California sea lions from coastal California. J Wildl Dis 2002, 38 (1), 7–17.

4. Prager, K. C.; Greig, D. J.; Alt, D. P.; Galloway, R. L.; Hornsby, R. L.; Palmer, L. J.; Soper, J.; Wu, Q.; Zuerner, R. L.; Gulland, F. M.; Lloyd-Smith, J. O., Asymptomatic and chronic carriage of Leptospira interrogans serovar Pomona in California sea lions (Zalophus californianus). Vet Microbiol 2013, 164 (1–2), 177–83.

5. Monahan, A. M.; Callanan, J. J.; Nally, J. E., Proteomic analysis of Leptospira interrogans shed in urine of chronically infected hosts. Infect Immun 2008, 76 (11), 4952–8.

6. Rojas, P.; Monahan, A. M.; Schuller, S.; Miller, I. S.; Markey, B. K.; Nally, J. E., Detection and quantification of leptospires in urine of dogs: a maintenance host for the zoonotic disease leptospirosis. Eur J Clin Microbiol Infect Dis 2010, 29 (10), 1305–9.

7. Eisner, C.; Faulhaber–Walter, R.; Wang, Y.; Leelahavanichkul, A.; Yuen, P. S. T.; Mizel, D.; Star, R. A.; Briggs, J. P.; Levine, M.; Schnermann, J., Major contribution of tubular secretion to creatinine clearance in mice. Kidney Int 2010, 77 (6), 519–526.

8. Sinkeler, S. J.; Visser, F. W.; Krikken, J. A.; Stegeman, C. A.; Homan van der Heide, J. J.; Navis, G., Higher body mass index is associated with higher fractional creatinine excretion in healthy subjects. Nephrology, dialysis, transplantation: official publication of the European Dialysis and Transplant Association – European Renal Association 2011, 26 (10), 3181–8.

9. Shemesh, O.; Golbetz, H.; Kriss, J. P.; Myers, B. D., Limitations of creatinine as a filtration marker in glomerulopathic patients. Kidney Int 1985, 28 (5), 830–8.

10. Doi, K.; Yuen, P. S.; Eisner, C.; Hu, X.; Leelahavanichkul, A.; Schnermann, J.; Star, R. A., Reduced production of creatinine limits its use as marker of kidney injury in sepsis. J Am Soc Nephrol 2009, 20 (6), 1217–21.

11. Perrone, R. D.; Madias, N. E.; Levey, A. S., Serum creatinine as an index of renal function: new insights into old concepts. Clin Chem 1992, 38 (10), 1933–53.

12. Moran, S. M.; Myers, B. D., Course of acute renal failure studied by a model of creatinine kinetics. Kidney Int 1985, 27 (6), 928–37.

13. Herget–Rosenthal, S.; Pietruck, F.; Volbracht, L.; Philipp, T.; Kribben, A., Serum cystatin C––a superior marker of rapidly reduced glomerular filtration after uninephrectomy in kidney donors compared to creatinine. Clin Nephrol 2005, 64 (1), 41–6.

14. Dennison, S. E.; Gulland, F. M.; Braselton, W. E., Standardized protocols for plasma clearance of iohexol are not appropriate for determination of glomerular filtration rates in anesthetized California sea lions (Zalophus californianus). J Zoo Wildl Med 2010, 41 (1), 144–7.

15. Hokamp, J. A.; Nabity, M. B., Renal biomarkers in domestic species. Veterinary clinical pathology 2016, 45 (1), 28–56.

16. Alge, J. L.; Arthur, J. M., Biomarkers of AKI: a review of mechanistic relevance and potential therapeutic implications. Clin J Am Soc Nephrol 2015, 10 (1), 147–55.

17. Wu, Q.; Prager, K. C.; Goldstein, T.; Alt, D. P.; Galloway, R. L.; Zuerner, R. L.; Lloyd–Smith, J. O.; Schwacke, L., Development of a real–time PCR for the detection of pathogenic Leptospira spp. in California sea lions. Dis Aquat Organ 2014, 110 (3), 165–172.

18. Dikken, H.; Kmety, E., Chapter VIII Serological Typing Methods of Leptospires. 1978, 11, 259–307.

19. Leon, I. R.; Schwammle, V.; Jensen, O. N.; Sprenger, R. R., Quantitative assessment of in–solution digestion efficiency identifies optimal protocols for unbiased protein analysis. Mol Cell Proteomics 2013, 12 (10), 2992–3005.

20. Vizcaino, J. A.; Csordas, A.; del–Toro, N.; Dianes, J. A.; Griss, J.; Lavidas, I.; Mayer, G.; Perez–Riverol, Y.; Reisinger, F.; Ternent, T.; Xu, Q. W.; Wang, R.; Hermjakob, H., 2016 update of the PRIDE database and its related tools. Nucleic Acids Res 2016, 44 (D1), D447–56.

21. Hochberg, Y.; Benjamini, Y., More powerful procedures for multiple significance testing. Stat Med 1990, 9 (7), 811–8.

22. Fabregat, A.; Jupe, S.; Matthews, L.; Sidiropoulos, K.; Gillespie, M.; Garapati, P.; Haw, R.; Jassal, B.; Korninger, F.; May, B.; Milacic, M.; Roca, C. D.; Rothfels, K.; Sevilla, C.; Shamovsky, V.; Shorser, S.; Varusai, T.; Viteri, G.; Weiser, J.; Wu, G.; Stein, L.; Hermjakob, H.; D’Eustachio, P., The Reactome Pathway Knowledgebase. Nucleic Acids Res 2018, 46 (D1), D649–D655.

23. Zhao, M.; Li, M.; Yang, Y.; Guo, Z.; Sun, Y.; Shao, C.; Sun, W.; Gao, Y., A comprehensive analysis and annotation of human normal urinary proteome. Sci Rep 2017, 7 (1), 3024.

24. Brandt, L. E.; Ehrhart, E. J.; Scherman, H.; Olver, C. S.; Bohn, A. A.; Prenni, J. E., Characterization of the canine urinary proteome. Veterinary clinical pathology 2014, 43 (2), 193–205.

25. Vaidya, V. S.; Ferguson, M. A.; Bonventre, J. V., Biomarkers of acute kidney injury. Annu Rev Pharmacol Toxicol 2008, 48, 463–93.

26. Nally, J. E.; Mullen, W.; Callanan, J. J.; Mischak, H.; Albalat, A., Detection of urinary biomarkers in reservoir hosts of leptospirosis by capillary electrophoresis–mass spectrometry. PROTEOMICS – Clinical Applications 2015, 9 (5–6), 543–551.

27. Ferlizza, E.; Campos, A.; Neagu, A.; Cuoghi, A.; Bellei, E.; Monari, E.; Dondi, F.; Almeida, A. M.; Isani, G., The effect of chronic kidney disease on the urine proteome in the domestic cat (Felis catus). Vet J 2015, 204 (1), 73–81.

28. Wenderfer, S. E.; Dubinsky, W. P.; Hernandez–Sanabria, M.; Braun, M. C., Urine proteome analysis in murine nephrotoxic serum nephritis. Am J Nephrol 2009, 30 (5), 450–8.

29. Lu, S. C.; Shieh, W. Y.; Chen, C. Y.; Hsu, S. C.; Chen, H. L., Lipopolysaccharide increases resistin gene expression in vivo and in vitro. FEBS Lett 2002, 530 (1–3), 158–62.

30. Ellison, R. T., 3rd; Giehl, T. J., Killing of gram–negative bacteria by lactoferrin and lysozyme. J Clin Invest 1991, 88 (4), 1080–91.

31. Zaragoza, C.; Barrera, R.; Centeno, F.; Tapia, J. A.; Mane, M. C., Characterization of renal damage in canine leptospirosis by sodium dodecyl sulphate–polyacrylamide gel electrophoresis (SDS–PAGE) and Western blotting of the urinary proteins. J Comp Pathol 2003, 129 (2–3), 169–78.

32. Phillips, J. A.; Holder, D. J.; Ennulat, D.; Gautier, J. C.; Sauer, J. M.; Yang, Y.; McDuffie, E.; Sonee, M.; Gu, Y. Z.; Troth, S. P.; Lynch, K.; Hamlin, D.; Peters, D. G.; Brees, D.; Walker, E. G., Rat Urinary Osteopontin and Neutrophil Gelatinase–Associated Lipocalin Improve Certainty of Detecting Drug– Induced Kidney Injury. Toxicol Sci 2016, 151 (2), 214–23.

33. Kashiwagi, E.; Tonomura, Y.; Kondo, C.; Masuno, K.; Fujisawa, K.; Tsuchiya, N.; Matsushima, S.; Torii, M.; Takasu, N.; Izawa, T.; Kuwamura, M.; Yamate, J., Involvement of neutrophil gelatinase– associated lipocalin and osteopontin in renal tubular regeneration and interstitial fibrosis after cisplatin– induced renal failure. Exp Toxicol Pathol 2014, 66 (7), 301–11.

34. Furuhashi, M.; Hotamisligil, G. S., Fatty acid–binding proteins: role in metabolic diseases and potential as drug targets. Nature reviews. Drug discovery 2008, 7 (6), 489.

35. Xu, Y.; Xie, Y.; Shao, X.; Ni, Z.; Mou, S., L–FABP: A novel biomarker of kidney disease. Clinica Chimica Acta 2015, 445, 85–90.

36. McMahon, B. A.; Murray, P. T., Urinary liver fatty acid–binding protein: another novel biomarker of acute kidney injury. Kidney Int 2010, 77 (8), 657–659.

37. Lindskog, C., The Human Protein Atlas – an important resource for basic and clinical research. Expert Rev Proteomics 2016, 13 (7), 627–9.

38. Garcia–Martinez, J. D.; Tvarijonaviciute, A.; Ceron, J. J.; Caldin, M.; Martinez–Subiela, S., Urinary clusterin as a renal marker in dogs. J Vet Diagn Invest 2012, 24 (2), 301–6.

39. Wagoner, M. P.; Yang, Y.; McDuffie, J. E.; Klapczynski, M.; Buck, W.; Cheatham, L.; Eisinger, D.; Sace, F.; Lynch, K. M.; Sonee, M.; Ma, J. Y.; Chen, Y.; Marshall, K.; Damour, M.; Stephen, L.; Dragan, Y. P.; Fikes, J.; Snook, S.; Kinter, L. B., Evaluation of Temporal Changes in Urine–based Metabolomic and Kidney Injury Markers to Detect Compound Induced Acute Kidney Tubular Toxicity in Beagle Dogs. Curr Top Med Chem 2017, 17 (24), 2767–2780.

40. Garimella, P. S.; Sarnak, M. J., Uromodulin in kidney health and disease. Curr Opin Nephrol Hypertens 2017, 26 (2), 136–142.

41. Garimella, P. S.; Bartz, T. M.; Ix, J. H.; Chonchol, M.; Shlipak, M. G.; Devarajan, P.; Bennett, M. R.; Sarnak, M. J., Urinary Uromodulin and Risk of Urinary Tract Infections: The Cardiovascular Health Study. Am J Kidney Dis 2017, 69 (6), 744–751.

42. Raila, J.; Schweigert, F. J.; Kohn, B., Relationship between urinary Tamm–Horsfall protein excretion and renal function in dogs with naturally occurring renal disease. Veterinary clinical pathology 2014, 43 (2), 261–5.

43. Hyde, G. D.; Taylor, R. F.; Ashton, N.; Borland, S. J.; Wu, H. S.; Gilmore, A. P.; Canfield, A. E., Axl tyrosine kinase protects against tubulo–interstitial apoptosis and progression of renal failure in a murine model of chronic kidney disease and hyperphosphataemia. PLoS One 2014, 9 (7), e102096.

44. Nagai, K.; Arai, H.; Yanagita, M.; Matsubara, T.; Kanamori, H.; Nakano, T.; Iehara, N.; Fukatsu, A.; Kita, T.; Doi, T., Growth arrest–specific gene 6 is involved in glomerular hypertrophy in the early stage of diabetic nephropathy. J Biol Chem 2003, 278 (20), 18229–34.

45. Ochodnicky, P.; Lattenist, L.; Ahdi, M.; Kers, J.; Uil, M.; Claessen, N.; Leemans, J. C.; Florquin, S.; Meijers, J. C. M.; Gerdes, V. E. A.; Roelofs, J. J. T. H., Increased Circulating and Urinary Levels of Soluble TAM Receptors in Diabetic Nephropathy. Am J Pathol 187 (9), 1971–1983.

46. Fiebeler, A.; Park, J. K.; Muller, D. N.; Lindschau, C.; Mengel, M.; Merkel, S.; Banas, B.; Luft, F. C.; Haller, H., Growth arrest specific protein 6/Axl signaling in human inflammatory renal diseases. Am J Kidney Dis 2004, 43 (2), 286–95.

47. White, F. N.; Ikeda, M.; Elsner, R. W., Adrenergic innervation of large arteries in the seal. Comp Gen Pharmacol 1973, 4 (15), 271–6.

48. Giangola, M. D.; Yang, W. L.; Rajayer, S. R.; Kuncewitch, M.; Molmenti, E.; Nicastro, J.; Coppa, G. F.; Wang, P., Growth arrest–specific protein 6 protects against renal ischemia–reperfusion injury. J Surg Res 2015, 199 (2), 572–9.

49. Korshunov, Vyacheslav A., Axl–dependent signalling: a clinical update. Clin Sci (Lond) 2012, 122 (8), 361–368.

50. Nally, J. E.; Monahan, A. M.; Miller, I. S.; Bonilla–Santiago, R.; Souda, P.; Whitelegge, J. P., Comparative proteomic analysis of differentially expressed proteins in the urine of reservoir hosts of leptospirosis. PLoS One 2011, 6 (10), e26046.

51. Velez, J. C., The importance of the intrarenal renin–angiotensin system. Nat Clin Pract Nephrol 2009, 5 (2), 89–100.

52. Morris, B. J.; Reid, I. A., A “Renin–Like” Enzymatic Action of Cathepsin D and the Similarity in Subcellular Distributions of “Renin–Like” Activity and Cathepsin D in the Midbrain of Dogs*. Endocrinology 1978, 103 (4), 1289–1296.

53. Pajenda, S.; Mechtler, K.; Wagner, L., Urinary neprilysin in the critically ill patient. BMC Nephrol 2017, 18, 172.

54. Chodavarapu, H.; Bradshaw, M.; Salem, E.; Elased, K., Decreased Renal and Urinary Neprilysin Protein Expression in db/db Mice. The FASEB Journal 2012, 26 (1_supplement), 1051.5–1051.5.

55. Yamaleyeva, L. M.; Gilliam–Davis, S.; Almeida, I.; Brosnihan, K. B.; Lindsey, S. H.; Chappell, M. C., Differential regulation of circulating and renal ACE2 and ACE in hypertensive mRen2.Lewis rats with early–onset diabetes. Am J Physiol Renal Physiol 2012, 302 (11), F1374–84.

56. Cesar, K. R.; Romero, E. C.; de Bragança, A. C.; Blanco, R. M.; Abreu, P. A. E.; Magaldi, A. J., Renal Involvement in Leptospirosis: The Effect of Glycolipoprotein on Renal Water Absorption. PLoS One. 2012, 7 (6), e37625. DOI:10.1371/journal.pone.0037625.

57. Etish, J. L.; Chapman, P. S.; Klag, A. R., Acquired nephrogenic diabetes insipidus in a dog with leptospirosis. Ir Vet J 2014, 67 (1), 7.

58. Magaldi, A. J.; Yasuda, P. N.; Kudo, L. H.; Seguro, A. C.; Rocha, A. S., Renal involvement in leptospirosis: a pathophysiologic study. Nephron 1992, 62 (3), 332–9.

59. Andersen, L. J.; Andersen, J. L.; Pump, B.; Bie, P., Natriuresis induced by mild hypernatremia in humans. American Journal of Physiology–Regulatory, Integrative and Comparative Physiology 2002, 282 (6), R1754–R1761.

60. Potthast, R.; Ehler, E.; Scheving, L. A.; Sindic, A.; Schlatter, E.; Kuhn, M., High Salt Intake Increases Uroguanyl in Expression in Mouse Kidney. Endocrinology 2001, 142 (7), 3087–3097.

61. Preston, R. A.; Afshartous, D.; Forte, L. R.; Rodco, R.; Alonso, A. B.; Garg, D.; Raij, L., Sodium challenge does not support an acute gastrointestinal–renal natriuretic signaling axis in humans. Kidney Int 2012, 82 (12), 1313–20.

